# Cluster Buster: A Machine Learning Algorithm for Genotyping SNPs from Raw Data

**DOI:** 10.1101/2024.08.23.609429

**Authors:** Jessica Martin, Nicole Kuznetsov, Kristin Levine, Mathew J. Koretsky, Samantha Hong, Mike A. Nalls, Dan Vitale

**Affiliations:** Center for Alzheimer’s and Related Dementias, National Institutes of Health, Bethesda, MD, USA 20892; DataTecnica LLC, Washington, DC, USA 20037

**Keywords:** Alzheimer’s disease, Parksinson’s disease, single nucleotide polymorphism, genome-wide association studies, GWAS, genetics, genotyping, prediction, neural network, machine learning

## Abstract

Genotyping single nucleotide polymorphisms (SNPs) is fundamental to disease research, as researchers seek to establish links between genetic variation and disease. Although significant advances in genome technology have been made with the development of bead-based SNP genotyping and Genome Studio software, some SNPs still fail to be genotyped, resulting in “no-calls” that impede downstream analyses. To recover these genotypes, we introduce Cluster Buster, a genotyping neural network and visual inspection system designed to improve the quality of neurodegenerative disease (NDD) research. Concordance analysis with whole genome sequencing (WGS) and imputed genotypes validated the reliability of predicted genotypes, with dozens of high-performing SNPs across *LRRK2, APOE*, and *GBA* loci achieving at least 90% concordance per SNP location. Further analysis of concordance between Genome Studio genotypes and imputed and WGS genotypes revealed discrepancies between the genotyping technologies, highlighting the need for selective application of Cluster Buster on SNP locations based on concordance rates. Cluster Buster’s implementation significantly reduces manual labor for recovering no-call SNPs, refining genotype quality for the Global Parkinson’s Genetics Program (GP2). This system facilitates better imputation and GWAS outcomes, ultimately contributing to a deeper understanding of genetic factors in NDDs.

## Introduction

In the last twenty years, genome-wide association studies (GWAS) have become the key to uncovering the statistical relationships between genetic variations and disease. They focus on single nucleotide polymorphisms (SNPs), which are variations in the genome capable of altering gene function and affecting disease heritability (Shastry, 2009). To power these studies, micro-array based technologies are widely used for genotyping SNPs. The NeuroBooster array is a genotyping platform focusing on SNPs associated with NDDs (Sara Bandres Ciga, 2023). These SNPs are often genotyped with Illumina’s bead-based SNP genotyping technology and GenomeStudio software. For some samples, specific SNPs fail to be genotyped by this algorithm and become “no-calls.” This missing data hampers genotype imputation, which relies on accurate genotype information to predict variants with linkage disequilibrium. This is critical for identifying genetic risk factors for disease. Consequently, this diminishes the quality of downstream analyses like GWAs that require high-resolution genetic data. Previously, these SNPs with missing genotypes had to be manually corrected by visual inspection, a laborious, time-intensive process (Sara Bandres Ciga, 2023).

Contributing accurately genotyped samples to the NeuroBooster array or other related technologies will help to improve genotype imputation quality in disease-related loci and, therefore, improve the power of GWAS studies on these SNPs. There is a need for a genotyping algorithm that can be used broadly on data submitted to the Global Parkinson’s Genetics Program (GP2) and other genetic cohort studies to greatly increase call rate for this purpose. To address this need, we introduce Cluster Buster, a genotyping neural network and visual inspection system that provides an efficient pipeline for genotyping missing variants. Here, we showcase the use of Cluster Buster in four NDD-related genes: *APOE, GBA, LRRK2*, and *SNCA*. This system vastly reduces the typical manual labor required to recover no-call SNPs in dozens of locations.

## Methods

### Data Collection and Preprocessing

#### GP2 Dataset

Each sample processed by GP2 was genotyped by the Illumina NeuroBooster Array SNP genotyping platform (Sara Bandres-Ciga, 2023). Primary sample identifiers, SNP genotypes, and metrics are collected and stored in parquet files. To minimize bias and ensure the representation of rare variants, 44 samples were randomly selected per the eleven available ancestries (Afro-Caribbean, African, Ashkenazi, Admixed American, Complex Admixture History, Central Asian, East Asian, European, Finish, Middle East, South Asian) (Dan Vitale, 2024). SNPs were selected to include those within *APOE, GBA, LRRK2*, and *SNCA*, for a total of 1,064 SNPs across 484 individuals directly genotyped on the NeuroBooster array.

#### Data Preprocessing

SNPs without a genotype cluster quality score from Illumina (GenTrain Score), a normalized intensity value (R), the ratio of allelic intensities (Theta), or genotype were filtered (Illumina, 2010). The genotype (GT) and single nucleotide polymorphism identifying string (snpID) were each converted from nominal into numerical categorical variables. Samples with an originally predicted genotype of “NC” (no-call) were separated and saved elsewhere (15,692 total SNPs). The remaining data were randomly divided into training (90%; 442,240 total SNPs), validation (5%; 24,570 total SNPs), and testing datasets (5%; 24,570 total SNPs).

#### Development of Genotyping Neural Network

##### Neural Network Architecture and Training

R values, Theta values, and snpIDs were selected as features for the neural network. The neural network was constructed with Tensorflow (version 2.16.1, Martín Abadi, 2015). Using the training dataset, the snpID variable was fed into an embedding layer that transforms the categorical variable into an information-dense vector (embedding size of 50) and then concatenated with the R and Theta numerical features. The concatenated features were fed into a dense layer with ReLu activation, followed by another dense layer with softmax activation. The targets were the one-hot encoded genotype calls from Illumina. Layer dimensions and hyperparameters were optimized using KerasTuner (version 1.4.6; O’Malley, 2015). The grid search tested layer sizes from 32 to 160 with a step size of 32. A callback (Keras EarlyStopping) was implemented with patience of 5 epochs with attention to validation loss. The Keras CategoricalFocalCrossentropy loss function and Adam optimizer function were used.

The final neural network architecture consisted of an input layer, an embedding layer for the snpID variable with an embedding size of 50 that was then flattened, a concatenation layer to concatenate the embedding output and the numerical features together, followed by a dense layer with 64 units, then a dense layer with 160 units, then a final output layer with 3 units (Figure 1B). This model was then trained with the training set for eleven epochs and a learning rate of 0.0001. The final accuracy on the training set was 0.9001 with a. loss of 2.3221E-5 and the final accuracy on the validation set was 0.9799 with a loss of 1.9199E-4. Predictions on the test set of SNPs and the set of SNPs with previous no-calls were then rendered with the trained genotyping neural network.

**Figure 1:**
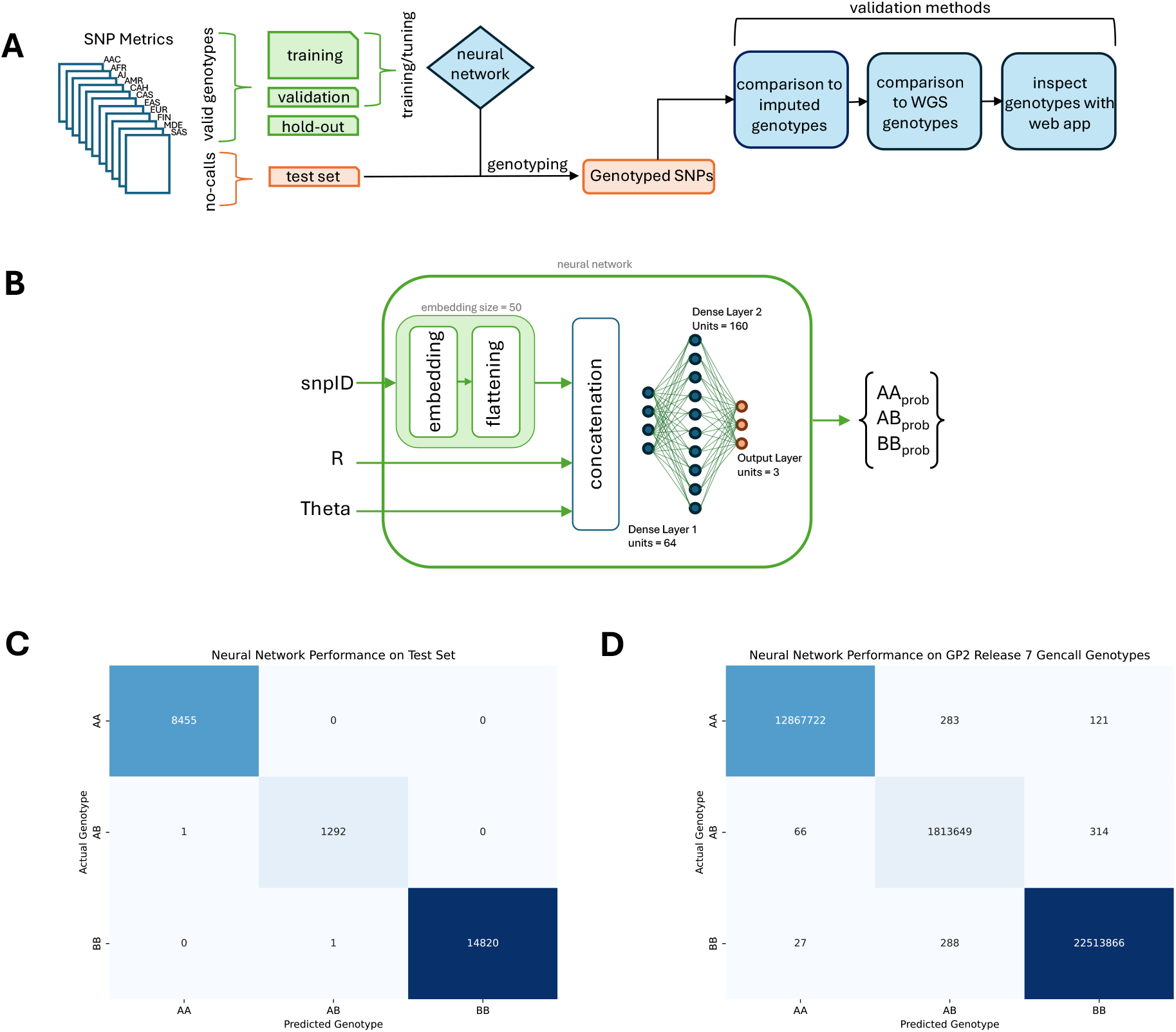
Cluster Buster Workflow and Performance. *A)* Flow of data through Cluster Buster. SNP metrics files from all available ancestries in GP2 are split into valid gencall (AA, AB, BB genotype) SNPs and no-call (NC genotype) SNPs. Valid genotypes are split 80-10-10 for training, validating, and testing the neural network. The trained neural network is applied to no-call SNPs. The predicted genotypes are compared to imputed and WGS genotypes and visually inspected with an online app. *B)* The structure of the neural network, including its inputs and outputs. *C)* Confusion matrix of neural network predicted genotypes versus gencall genotypes on the hold-out test set. *D)* Confusion matrix of neural network predicted genotypes versus gencall genotypes on the entirety of non-GDPR release7 samples from GP2.

##### Comparison of Predicted Genotypes to Imputed Genotypes

Samples from GP2 release 6 with imputation calls from the TOPMed server (Das et al., 2016) hosted on Biowulf were stored in PLINK2 format (pgen, psam, and pvar files) for each ancestry available (version v2.00a5.10LM, Shaun Purcell, Christopher Chang). Data was extracted for SNPs within the chromosome base pair ranges for each gene locus and extracted into PLINK2 .raw files. Samples were then matched on their GP2SampleID and matched per SNP on the matching chromosome, base position, reference allele, and alternate allele.

##### Comparison of Predicted Genotypes to WGS Genotypes

Samples from GP2 release 6 were sequenced and called with DeepVariant-GLnexus (Hampton Leonard et al., 2024). These samples with whole genome sequence genotype calls hosted on Terra were stored in PLINK2 format (pgen, psam, and pvar files) for each ancestry available. Extraction and matching were done the same as for imputed genotypes described previously.

Data was extracted for SNPs within the chromosome, base pair ranges for each gene locus and extracted into PLINK2 .raw files (version v2.00a5.10LM, Shaun Purcell, Christopher Chang). Samples were then matched with predicted genotypes by matching chromosomes, base position, reference allele, and alternate allele.

## Results

### Genotyping with the Neural Network

After training the genotyping neural network (Figure 1A and 1B), the neural network was applied to the hold-out set of SNPs to evaluate its performance on unseen SNPs using Illumina probe measurements R and Theta and a categorical variable representing snpID as inputs per SNP. The neural network genotyped the SNPs in this set with an accuracy of 99.9% (Figure 1C). Then, the genotyping neural network was applied to all SNPs in GP2 release 7 samples with valid (not no-call) genotypes rendered with the Illumina Gencall algorithm (the software used to genotype within GenomeStudio). For 36,592 samples at 1,064 different SNP locations for 37,196,336 SNPs, the neural network predicted genotypes that matched the Gencall genotypes at 99.9% (Figure 1D). The genotyping neural network was then applied to all SNPs across GP2 release 7 samples with no-calls to recover their genotypes (1,129,556 SNPs).

### Validation of Predicted Genotypes with Imputation and Whole Genome Sequencing

For samples in which imputation genotype calls and/or whole genome sequencing (WGS) genotype calls were available, predicted genotypes were compared to imputed and WGS genotypes. This analysis covered 569 SNP locations. Rates of concordance between predicted genotypes, imputed genotypes, and WGS were calculated on a per SNP level as a metric of predicted genotype reliability and can be found in Supplementary Table 1. For a SNP to be considered “high-performing” and used to reliably recover genotypes for GP2, the available concordance rates between predicted genotypes and imputed genotypes and predicted genotypes and WGS genotypes needed to be 90% or higher. 43 SNP locations in *LRRK2*, 30 SNP locations in *APOE*, and 28 SNP locations in *GBA* met this threshold. The recovered genotypes of SNPs in these locations can be included in imputation efforts and future GWAS. SNPs with predicted genotypes that matched either imputed or WGS genotypes were used to recalculate the overall call rate for the SNP location in GP2, which can be found in Supplementary Table 2. Examples of SNPs with high concordance with their predicted genotypes on allele measurement plots are shown in Figure 2. When plotting per-SNP measurements for R on the y-axis and Theta on the x-axis, genotypes generally follow a visual clustering behavior where genotypes homozygous for the major and minor allele fall on the left and right sides of the plot, respectfully, and heterozygous genotypes appear towards the middle of the plot. SNPs with predicted genotypes highly concordant with imputation and WGS genotypes generally followed that visual pattern.

**Figure 2:**
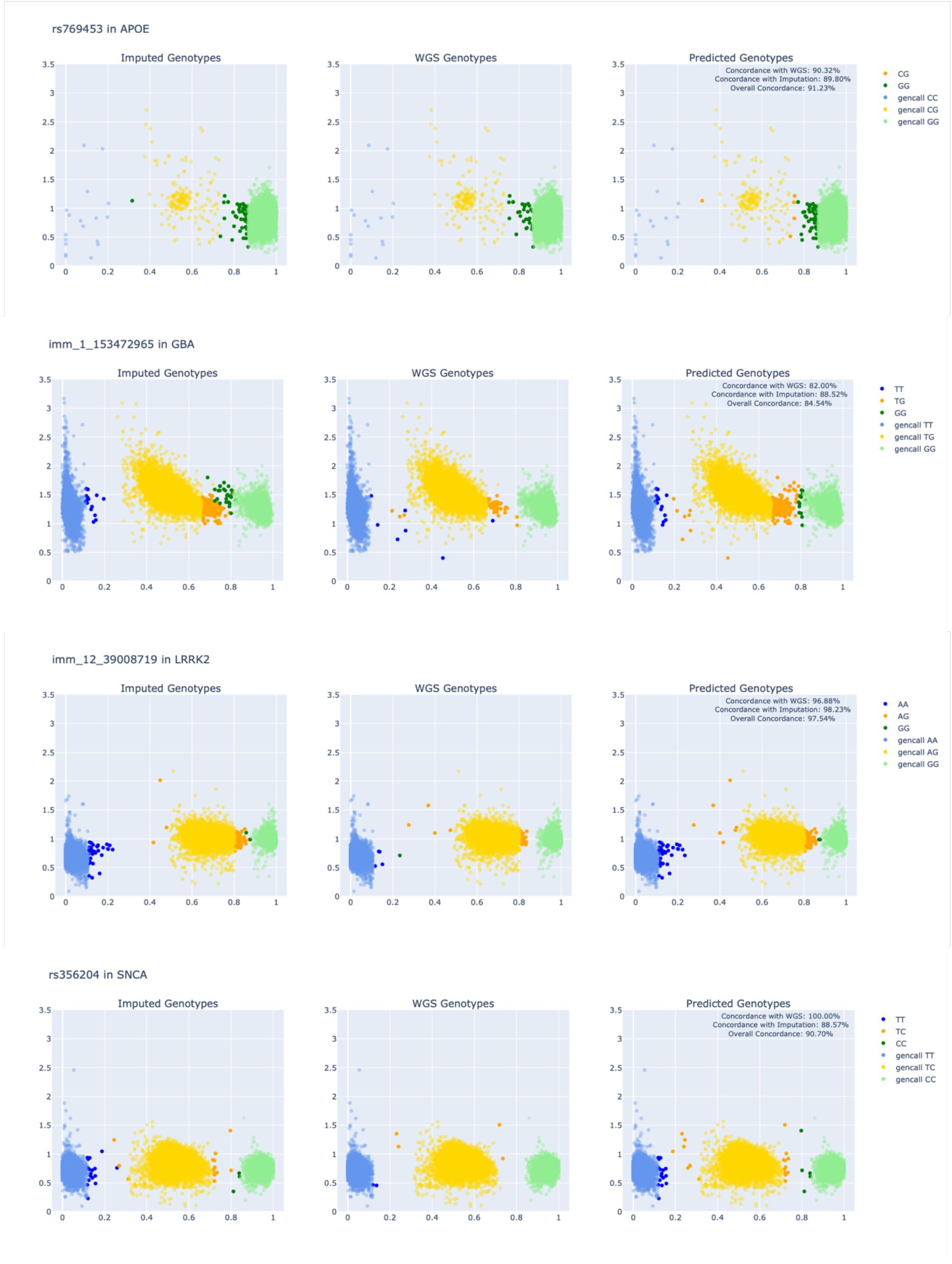
SNPs with High Concordance between Predicted Genotypes and Imputed, WGS genotypes. *A)* Three scatter plots showing imputed, WGS, and Cluster Buster predicted genotypes for a SNP in APOE for non-GDPR samples in GP2 release7. A transparent layer of the valid gencall genotypes for the SNP appear in the background of the scatter plots. *B)* Three scatter plots showing imputed, WGS, and Cluster Buster predicted genotypes for a SNP in GBA for non-GDPR samples in GP2 release7. A transparent layer of the valid gencall genotypes for the SNP appear in the background of the scatter plots. *C)* Three scatter plots showing imputed, WGS, and Cluster Buster predicted genotypes for a SNP in LRRK2 for non-GDPR samples in GP2 release7. A transparent layer of the valid gencall genotypes for the SNP appear in the background of the scatter plots. *D)* Three scatter plots showing imputed, WGS, and Cluster Buster predicted genotypes for a SNP in SNCA for non-GDPR samples in GP2 release7. A transparent layer of the valid gencall genotypes for the SNP appear in the background of the scatter plots.

Illumina Inc. records a GenTrain Score ranging from 0.00 to 1.00 for each SNP location, which statistically represents the degree to which genotypes for that SNP follow an “expected” clustering behavior (Illumina Inc, 2005). A higher GenTrain Score indicates that raw data for the SNP is more likely to cluster based on genotype. Pairwise Pearson correlation between a SNPs given GenTrain score and its overall concordance with imputation revealed only a modest positive relationship with a correlation coefficient of r=0.32. There was virtually no correlation between the GenTrain score and a SNP concordance rate with WGS (r=0.007 correlation coefficient). This suggests that Cluster Buster’s performance is largely independent of GenTrain Score and is suitable for application to SNPs with lower GenTrain Scores.

Exploring the SNPs with the most discordance between predicted genotypes and WGS and imputed genotypes revealed that for specific SNPs, the imputed and WGS genotypes don’t follow the expected pattern of genotype clustering when plotting allele measurements from Illumina raw data. Examples of poorly predicted SNP genotypes and their respective imputed and WGS genotypes can be seen in allele measurement plots in Figure 3. The canonical SNPs involved in APOE E2 and E4 alleles, rs7412 and rs429358 (Sweigart et al., 2021), had low rates of concordance between predicted genotypes and imputed and WGS genotypes for this reason.

**Figure 3:**
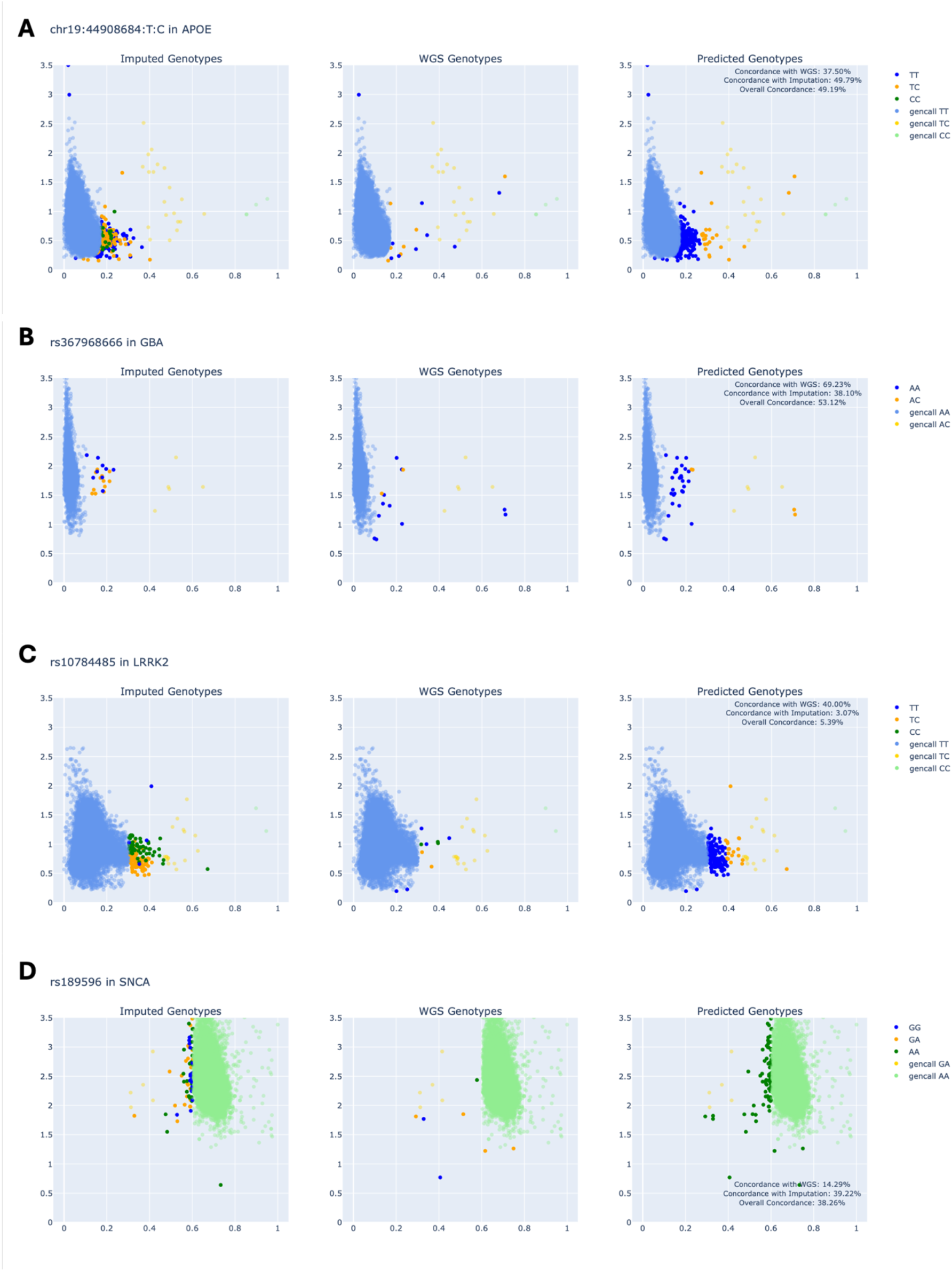
SNPs with Low Concordance between Predicted Genotypes and Imputed, WGS genotypes. *A)* Three scatter plots showing imputed, WGS, and Cluster Buster predicted genotypes for a SNP in APOE for non-GDPR samples in GP2 release7. A transparent layer of the valid gencall genotypes for the SNP appear in the background of the scatter plots. *B)* Three scatter plots showing imputed, WGS, and Cluster Buster predicted genotypes for a SNP in GBA for non-GDPR samples in GP2 release7. A transparent layer of the valid gencall genotypes for the SNP appear in the background of the scatter plots. *C)* Three scatter plots showing imputed, WGS, and Cluster Buster predicted genotypes for a SNP in LRRK2 for non-GDPR samples in GP2 release7. A transparent layer of the valid gencall genotypes for the SNP appear in the background of the scatter plots. *D)* Three scatter plots showing imputed, WGS, and Cluster Buster predicted genotypes for a SNP in SNCA for non-GDPR samples in GP2 release7. A transparent layer of the valid gencall genotypes for the SNP appear in the background of the scatter plots.

### Exploration of Gencall Genotypes and Concordance with Imputation and WGS

Finding SNPs that did not adhere to expected clustering behavior led to an analysis of concordance between valid genotypes from GenomeStudio and their imputed and WGS genotypes. Rates of concordance per SNP location can be found in Supplementary Table 3. 14 SNP locations in *LRRK2*, five SNP locations in *SNCA*, and three SNP locations in *APOE* had concordance rates below 90% either when comparing Gencall genotypes to WGS genotypes or when comparing Gencall genotypes to imputed genotypes. Notably, the SNP associated with the APOE E2 and E4 alleles (rs429358) had a Gencall genotype concordance rate with imputed genotypes of 78.9% and a Gencall genotype concordance rate with WGS genotypes of 68.8%. The other SNP associated with APOE E2 and E4 (rs7412) had better concordance rates: 90% concordance between Gencall and imputed genotypes and 85% concordance between Gencall and WGS genotypes. For these SNP locations with low concordance between Gencall genotypes and imputed or WGS genotypes, the only reliable way to genotype is to impute with haplotype information or use whole-genome sequencing technology.

GP2 monitors SNPs that are unsuitable for genotyping with the Gencall algorithm as part of the quality control process. Three SNP locations in *APOE*, 21 in *LRRK2*, and two in *SNCA* were not already tracked by GP2 and can be added to the list of SNPs to exclude from Gencall genotyping (Figure 4). Separating SNPs by GP2 quality-control status reveals that much of Cluster Buster’s poor performance on certain SNP locations can be attributed to those SNP locations not being reliably genotyped by the array, rendering the raw data used for predicting genotypes to be useless at these locations. SNPs on the exclude list exhibited lower concordance rates between predicted genotypes and summary concordance (concordance with either imputation or WGS) (Figure 5). Mann-Whitney U-tests revealed significant differences in summary concordance rates between included and excluded SNPs across three gene regions: *APOE* (p = 0.0005), *LRRK2* (p = 0.001), and *SNCA* (p = 0.002).

**Figure 4:**
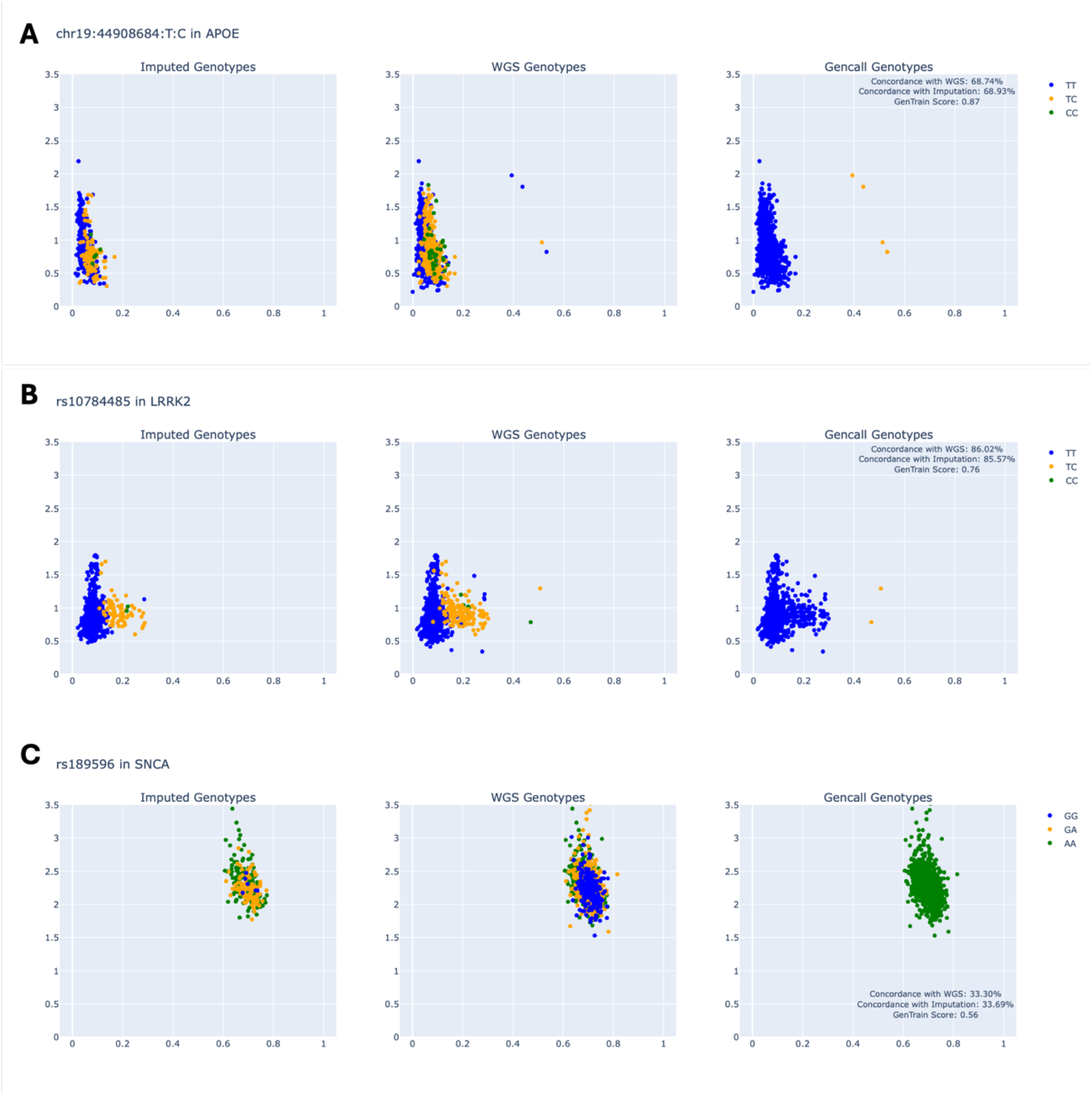
SNPs with Low Concordance between Gencall Genotypes and Imputed, WGS genotypes. *A)* Three scatter plots showing imputed, WGS, and gencall genotypes for a SNP in APOE for non-GDPR samples in GP2 release7. *B)* Three scatter plots showing imputed, WGS, and gencall genotypes for a SNP in LRRK2 for non-GDPR samples in GP2 release7. *C)* Three scatter plots showing imputed, WGS, and gencall genotypes for a SNP in SNCA for non-GDPR samples in GP2 release7.

**Figure 5.**
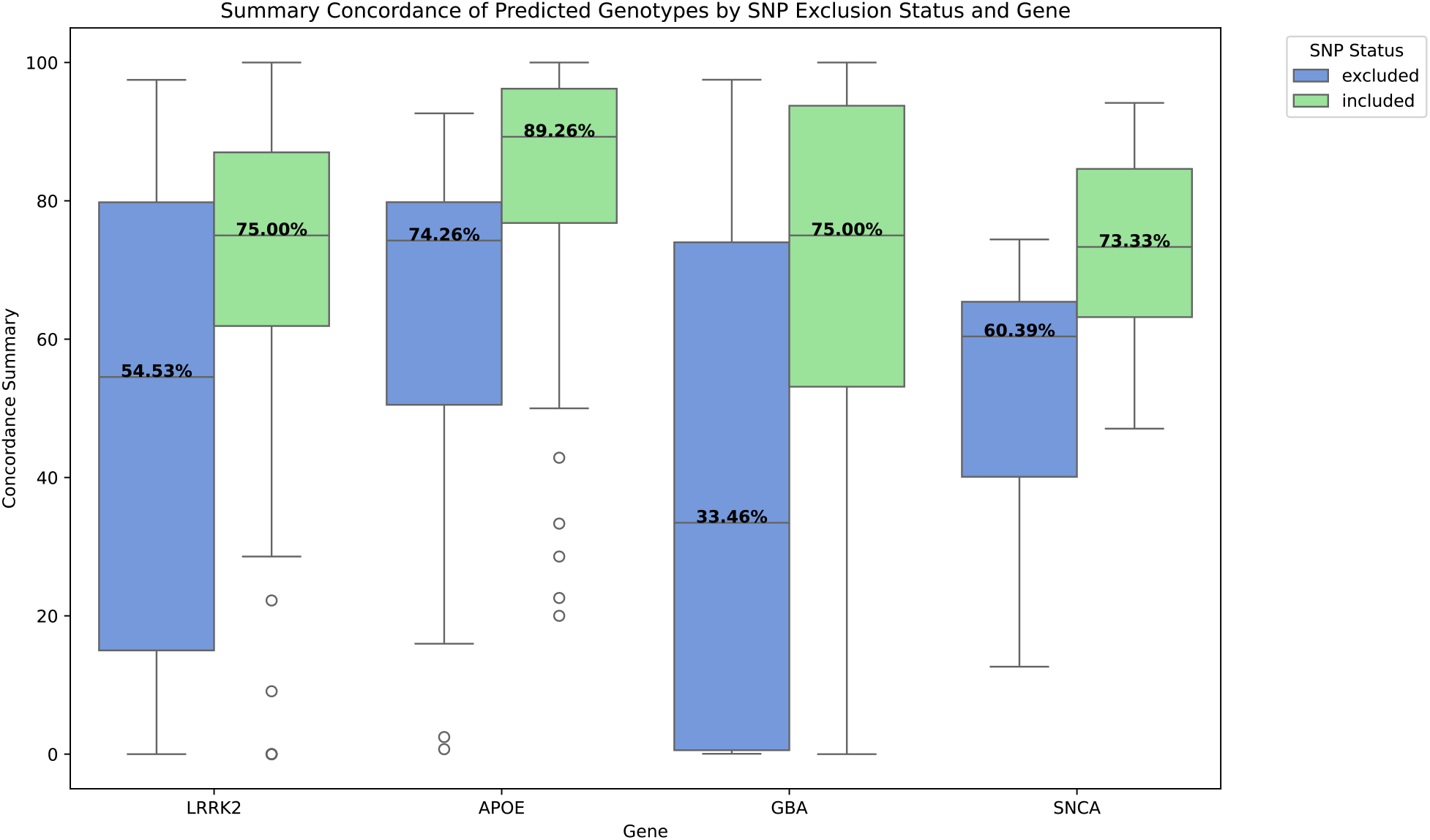
How SNP Status (Included or Excluded by GP2) Affects Cluster Buster Performance per Gene. Two boxplots per gene show how predicted genotypes from Cluster Buster have a higher concordance rate with imputed and WGS genotypes if they are SNPs trusted (SNP status - included) by GP2. Performance dips on SNPs that are excluded by GP2 during quality control.

## Discussion

By leveraging raw data from Illumina BeadStation technology, the Cluster Buster neural network demonstrated exceptional accuracy in genotype predictions for dozens of SNPs across a large dataset. This approach not only facilitated the recovery of missing genotypes but also provided a new avenue for SNP analysis in regions where more traditional genotyping technologies falter.

For specific SNPs in all four studied gene loci, both imputed and WGS genotypes contradicted the expected clustering behavior based on R and Theta probe measurements. The genotypes rendered for these SNPs by Illumina Gencall software may not be trustworthy, and therefore, the genotyping may be better left to imputation and WGS techniques. Because of this finding, determining which SNPs can be genotyped accurately with Cluster Buster requires selectivity based on concordance with other genotyping methods.

Analysis of SNPs with low concordance rates between predicted, imputed, and WGS genotypes uncovered inconsistencies between genotyping technologies, leading to analysis of the concordance rates between previously accepted Gencall genotypes and their respective imputed and WGS genotypes. This investigation revealed discrepancies for certain SNPs that point to potential instrumentation errors. However, neither imputation nor WGS are foolproof technologies, and it must be noted that concordance rates of genotypes between technologies offer only an approximation of the accuracy of the predicted genotype.

In part, GP2’s quality control pipeline monitors SNPs that require imputation or WGS to genotype rather than Illumina’s SNP genotyping platform. This detailed exploration of genotype concordance between technologies adds 26 SNP locations to GP2’s efforts to improve the overall accuracy and reliability of genotyping data for its cohorts.

As part of ongoing research, a method is being developed to flag which SNPs do not exhibit genotype clustering behavior according to probe measurements. These SNPs therefore may not be appropriate for the application of Cluster Buster. The centroid and width of each genotype cluster can be calculated using imputed genotypes and R and Theta values from Illumina for each SNP. Then, a SNP may be flagged for non-clustering behavior if genotype clusters are too wide, the genotype cluster centroids are too close, or the genotype centroids have very low R-value coordinates. The specific thresholds for delineating these conditions are currently being explored. Developing this method will allow researchers to automatically determine which SNPs are inappropriate for Illumina Gencall data analysis software or Cluster Buster without visual inspection.

Future research will center on expanding Cluster Buster’s capabilities to genotype more SNP locations in the NeuroBooster array. This will require training the neural network in a broader range of SNPs and acquiring more imputation and WGS genotyping for validation.

## Conclusions

Cluster Buster rapidly recovers SNP genotypes using raw data from cost-effective bead-based SNP genotyping technology. The neural network has been carefully trained on ancestrally diverse data and variants of all rarities to ensure minimal bias and maximum applicability.

Currently, the genotyping system verifiably genotypes 43 SNP locations in *LRRK2*, 30 SNP locations in *APOE*, and 28 SNP locations in *GBA* at a concordance rate of 90% or better with genotypes from TOPMed server and whole genome sequencing. Exploration of concordance rates indicated several SNPs that do not exhibit genotype clustering behavior based on probe measurements from Illumina GenomeStudio software, suggesting that these particular SNPs are better analyzed using whole genome sequencing. Cluster Buster provides a scalable, efficient solution for improving genotype data quality in biobank-scale analysis. Genotypes recovered with Cluster Buster will improve imputation efforts on diverse populations and increase GWAS power in future studies.

## Acknowledgments

This research was supported in part by the Intramural Research Program of the NIH, National Institute on Aging (NIA), National Institutes of Health, Department of Health and Human Services, project number ZO1 AG000534, and the National Institute of Neurological Disorders and Stroke.

Data (DOI 10.5281/zenodo.10962119, release 7) used in the preparation of this article were obtained from the Global Parkinson’s Genetics Program (GP2). GP2 is funded by the Aligning Science Against Parkinson’s (ASAP) Initiative and implemented by The Michael J. Fox Foundation for Parkinson’s Research (https://www.gp2.org). For a complete list of GP2 members, see http://www.gp2.org.

Data used in the preparation of this article were obtained from the Accelerating Medicine Partnership® (AMP®) Parkinson’s Disease (AMP PD) Knowledge Platform. For up-to-date information on the study, visit https://www.amp-pd.org. The AMP® PD program is a public-private partnership managed by the Foundation for the National Institutes of Health and funded by the National Institute of Neurological Disorders and Stroke (NINDS) in partnership with the Aligning Science Across Parkinson’s (ASAP) initiative; Celgene Corporation, a subsidiary of Bristol-Myers Squibb Company; GlaxoSmithKline plc (GSK); The Michael J. Fox Foundation for Parkinson’s Research; Pfizer Inc.; AbbVie Inc.; Sanofi US Services Inc.; and Verily Life Sciences. ACCELERATING MEDICINES PARTNERSHIP and AMP are registered service marks of the U.S. Department of Health and Human Services.

